## Supplementary Material for "The influence of diet on gut microbiome and body mass dynamics in a capital-breeding migratory bird"

**Foraging phenotype modifies the gut microbiome and alters body mass gain of a capital breeder.**

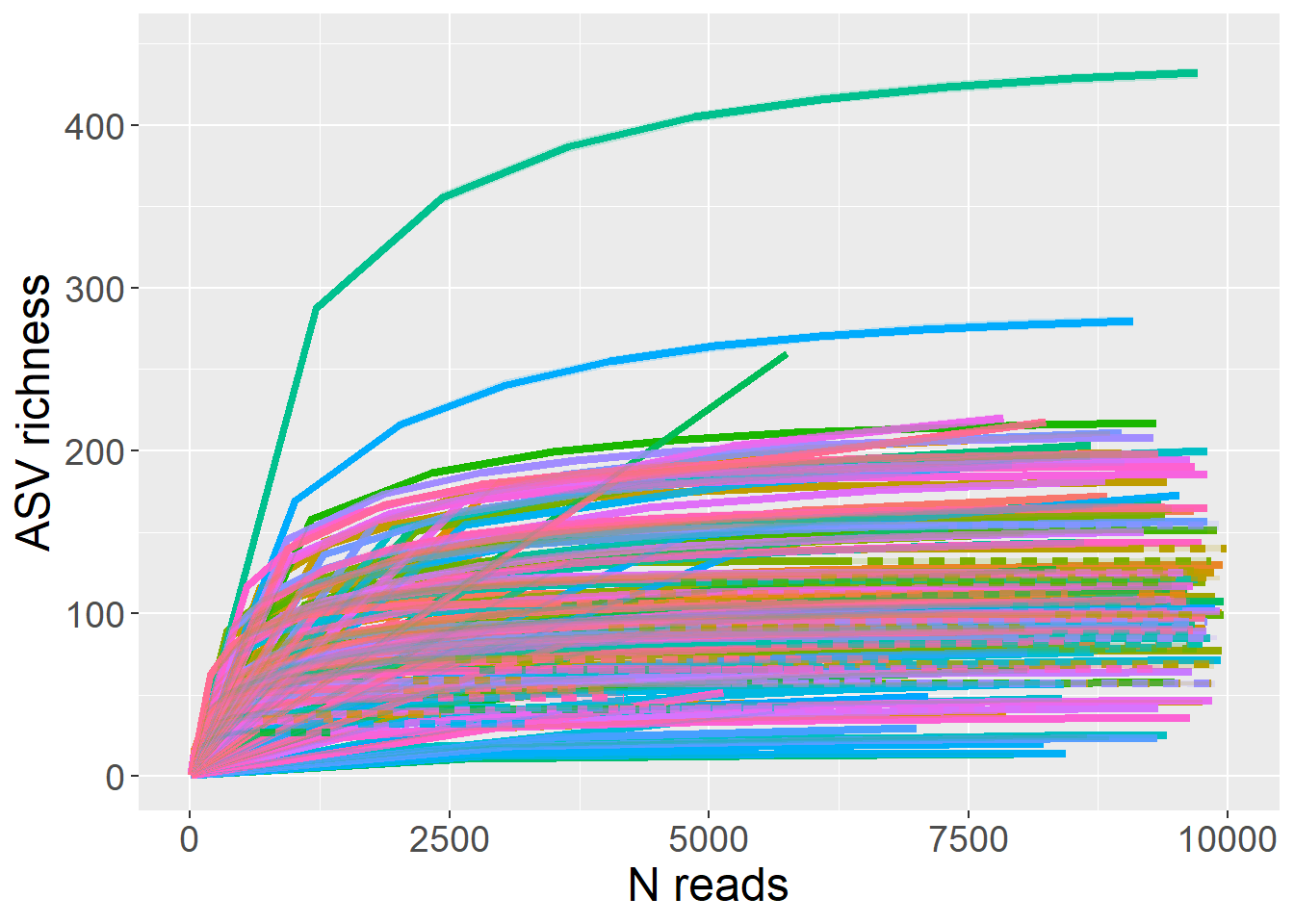

**Figure S1:** Rarefaction curves to estimate the ASV richness expected from samples at different sampling depths, generated in the R package iNEXT. Curves for each sample (n=145) are shown after filtering of contaminants.

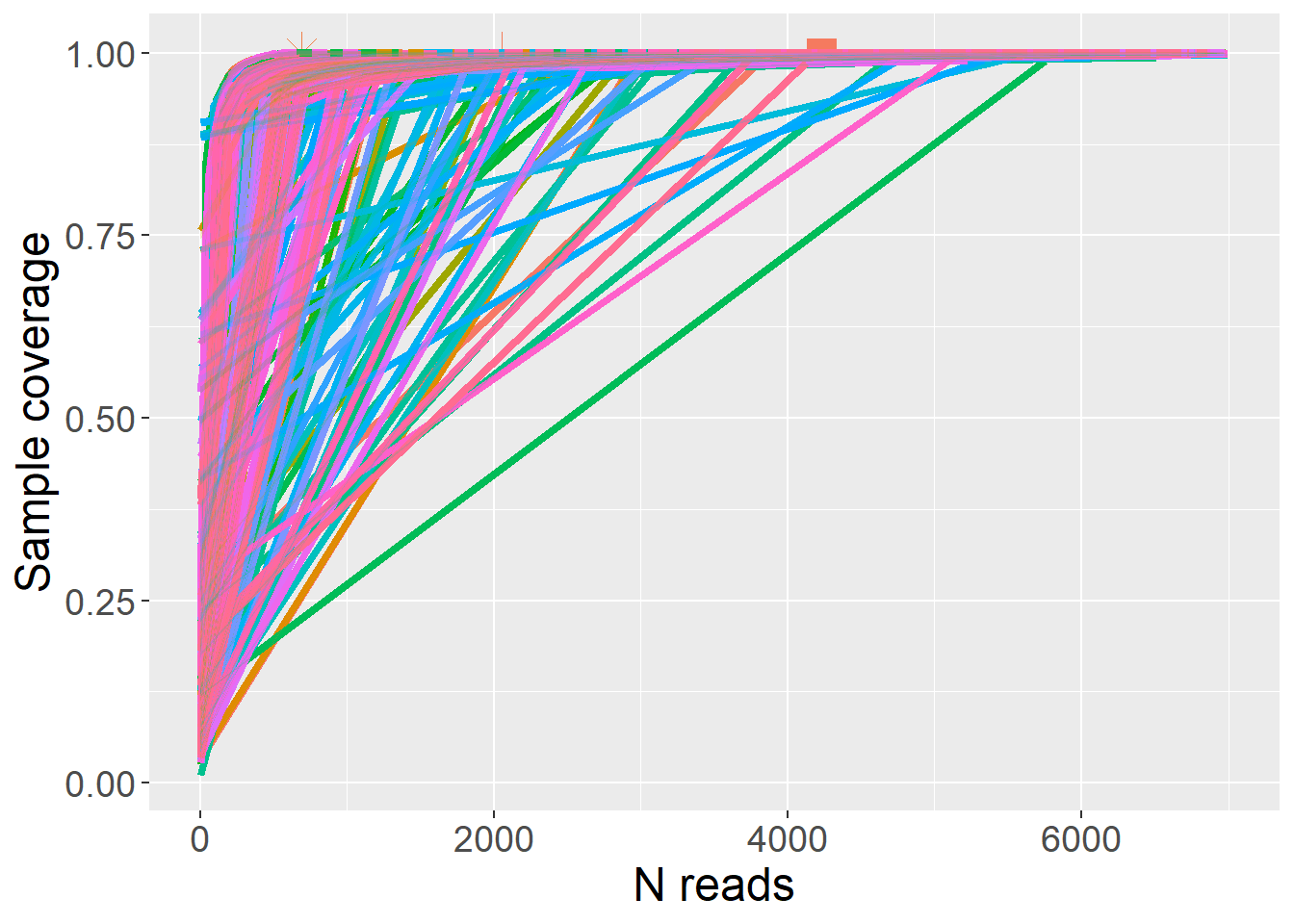

**Figure S2:** Sample coverage curves to estimate the coverage of samples at different sampling depths, generated in the R package iNEXT. Curves for each sample (n=145) are shown after filtering of contaminants.

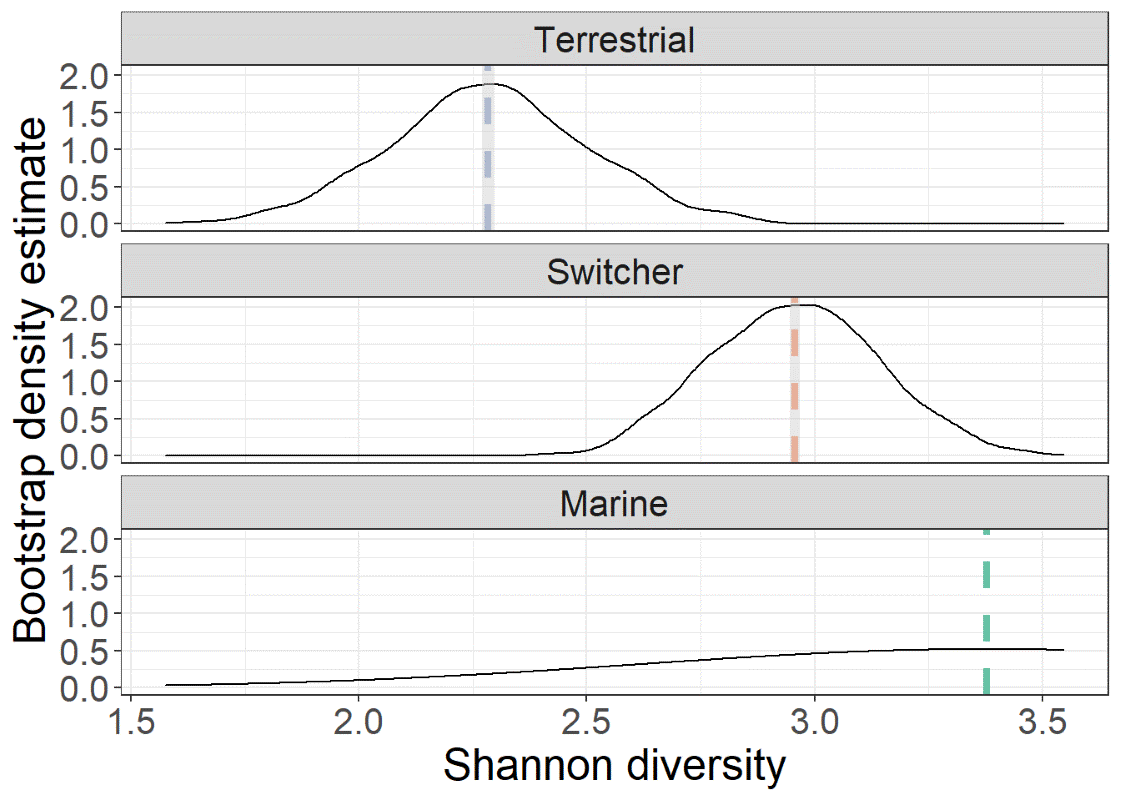

**Figure S3:** Bootstrap density estimates of Shannon diversity for each foraging phenotype. The rarified microbiome data was subsampled to the minimum group size for foraging phenotype (n=16, marine specialists) and the average Shannon diversity value was calculated across 1000 iterations to produce a distribution of bootstrapped Shannon diversity estimates.

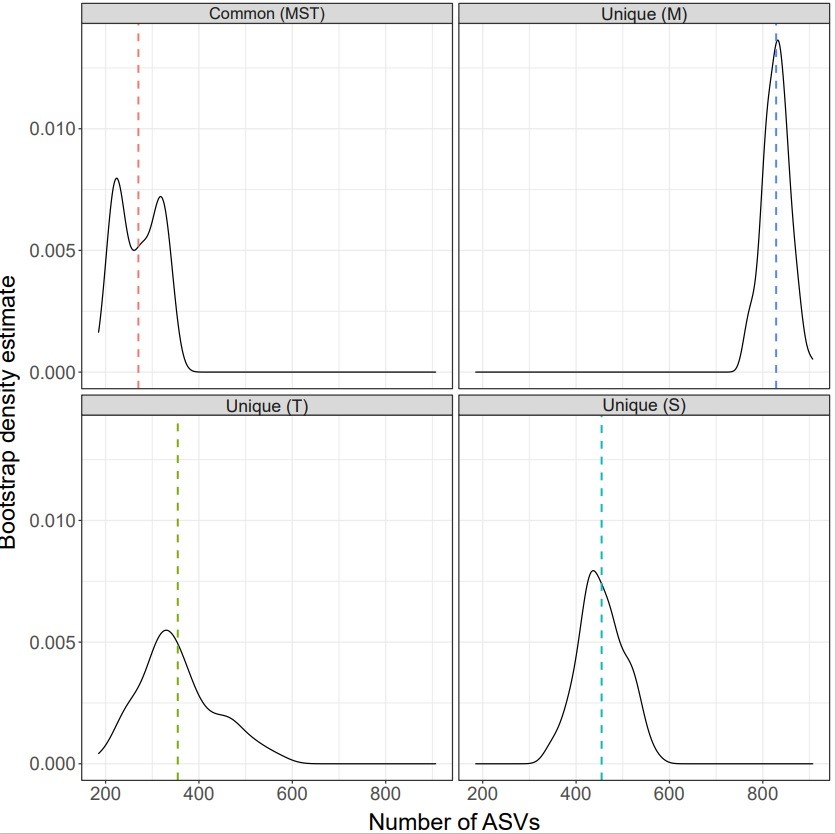

**Figure S4:** Faecal microbial communities (unrarefied) sampled from brent geese were categorised into groups based on the foraging behaviour of the individuals. To account for the uneven sampling effort between these groups, bootstrap subsampling to the minimum sample size (n=17, marine group) was performed with 1000 iterations to calculate the number of unique and shared bacterial amplicon sequence variants (ASVs) among foraging phenotypes (M=marine, T=terrestrial, S=switcher). The densities of these bootstrap estimates are shown, with the mean number of ASVs across iterations indicated with a dashed line.

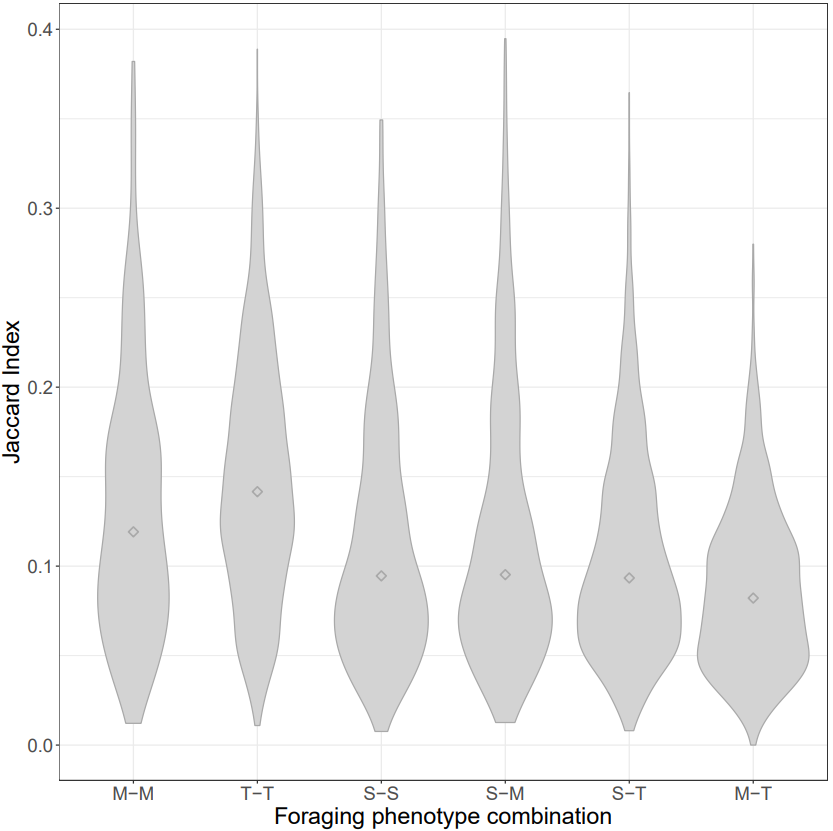

**Figure S5:** Proportions of shared microbial amplicon sequence variants (ASVs) between foraging phenotypes of light-bellied brent geese. Faecal microbial communities were categorised based on the foraging phenotype of the individual host; marine (M, n=17), terrestrial (T, n=57), and individuals switching between these two habitats (S, n=34). The Jaccard Index was used to show the proportion of ASVs shared between pairs of faecal samples from the same or different foraging phenotypes. A violin plot was used to show the distribution of Jaccard Index values, with the mean per comparison indicated by a diamond shape.

**Table S1:** Indicator analysis (R package *labdsv*) displaying the ASVs identified as indicative of foraging preference at the genus level.

| **ASV** | **Foraging Preference** | **P value** | | **Family** | **Genus** | |
| --- | --- | --- | --- | --- | --- | --- |
| SV2 | Terrestrial | | 0.003 | *Pseudomonadaceae* | | *Pseudomonas* |
| SV3 | Terrestrial | | 0.020 | *Pseudomonadaceae* | | *Pseudomonas* |
| SV6 | Terrestrial | | 0.043 | *Lactobacillaceae* | | *Lactobacillus* |
| SV28 | Marine | | 0.001 | *Phyllobacteriaceae* | | *Hoeflea* |
| SV31 | Terrestrial | | 0.001 | *Rhizobiaceae* | | *Rhizobium* |
| SV36 | Marine | | 0.001 | *Hyphomicrobiaceae* | | *Devosia* |

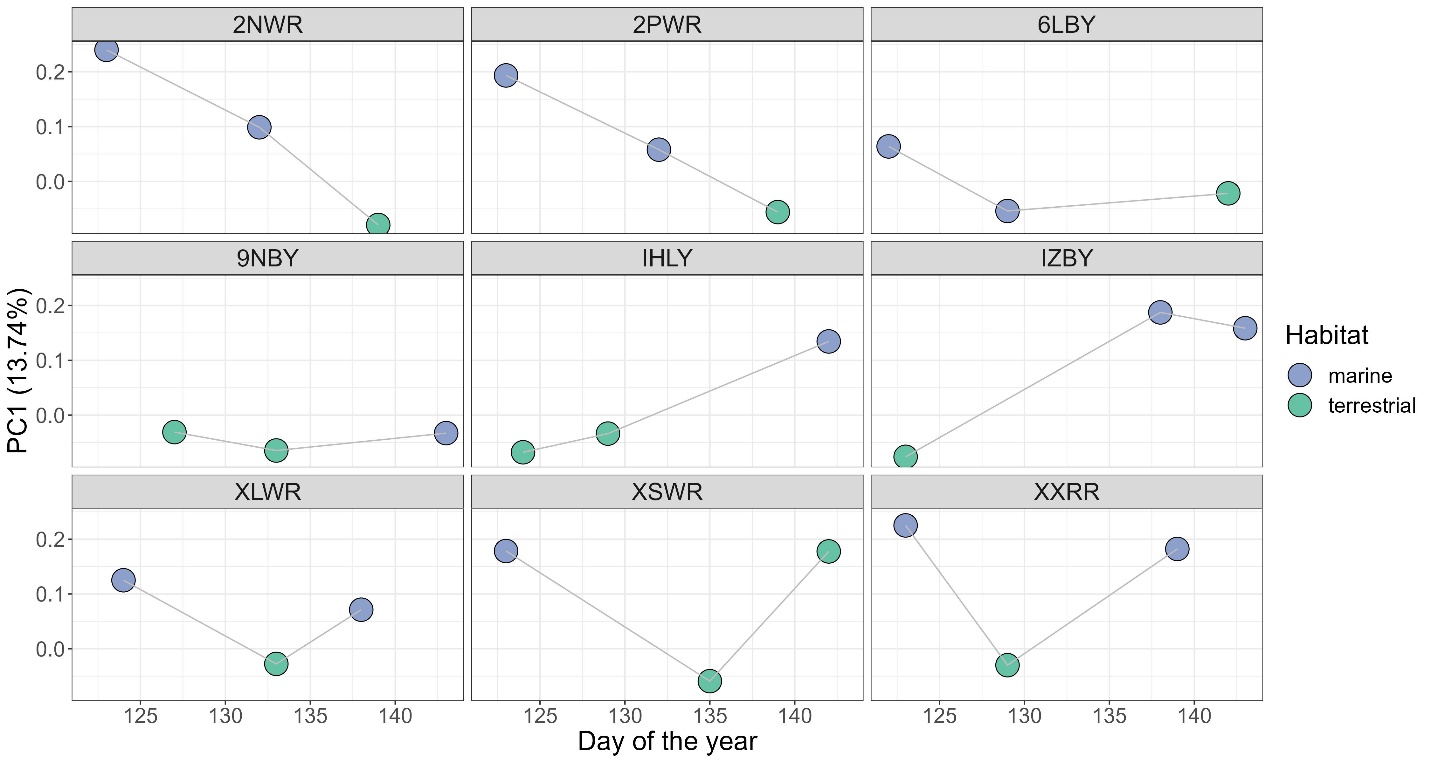

**Figure S6:** Microbiome compositional change over time for individual switchers. Changes in the position along the first axis of a Principle Component Analysis (PC1) over time are shown for individual geese that were sampled across different habitat types; marine (blue) and terrestrial (green). The alphanumeric codes at the top of each panel refer to the individual switcher (geese that moved among different habitat types, n=9). Colours indicate the habitat type in use at the point of faecal sample collection.

**Table S2:** Linear mixed model selection for body condition (API) variation, within delta six AICc units of the best model. Model number, foraging phenotype, sex, standardised day, quadratic effect of standardised day (Day^2^), the interaction of foraging phenotype and standardised day (Feeder Type:z_Day), the interaction of foraging phenotype and the quadratic effect of standardised day (Feeder Type:Day^2^), AICc score and models retained according to the nesting rule.

| **Model number** | **Foraging phenotype** | **Sex** | **Day** | **Day^2^** | **Foraging phenotype: Day** | **Foraging phenotype: Day^2^** | **AICc** | **Retained** |
| --- | --- | --- | --- | --- | --- | --- | --- | --- |
| 1 |  |  | 1.1030 |  |  |  | 414.2 |  |
| 2 | + | ­+ | 1.0970 | 0.03322 |  |  | 415.2 | / |
| 3 | + | + | 1.0290 | 0.04742 | + |  | 416.4 |  |
| 4 | + | + | 1.1040 |  |  |  | 416.4 |  |
| 5 | + | + | 1.0550 |  | + |  | 416.5 |  |
| 6 | + | + | 1.0980 | 0.03258 |  |  | 417.5 |  |
| 7 | + | + | 0.9588 | 0.17800 | + | + | 418.0 |  |
